# Bee visitation and fruit quality in berries under protected cropping vary along the length of polytunnels

**DOI:** 10.1101/722041

**Authors:** Mark Hall, Jeremy Jones, Maurizio Rocchetti, Derek Wright, Romina Rader

**Affiliations:** School of Environmental and Rural Science, University of New England, Armidale, NSW 2350, Australia; Costagroup, Corindi Berry Farm, Corindi NSW 2456, Australia; Hawkesbury Institute for the Environment, Western Sydney University, Richmond, NSW 2753, Australia

**Author notes:** Co-lead authors (Mark Hall, Jeremy Jones).

**Keywords:** *Apis mellifera*, blueberry, *Homalictus urbanus*, pollinator, raspberry, *Tetragonula carbonaria*

## Abstract

Wild and managed bees provide effective crop pollination services worldwide. Protected cropping conditions are thought to alter the ambient environmental conditions in which pollinators forage for flowers, yet few studies have compared conditions at the edges and centre of growing tunnels. We measured environmental variables (temperature, relative humidity, wind speed, white light and UV light) and surveyed the activity of managed honeybees *Apis mellifera*, wild stingless bees *Tetragonula carbonaria* and sweat bees *Homalictus urbanus* along the length of 32 multiple open-ended polyethylene growing tunnels. These were spaced across 12 blocks at two commercial berry farms, in Coffs Harbour, New South Wales and Walkamin, North Queensland, Australia. Berry yield, fresh weight and other quality metrics were recorded at discrete increments along the length of the tunnels. We found a higher abundance and greater number of flower visits by stingless bees and honeybees at the end of tunnels, and less frequent visits to flowers toward the middle of tunnels. The centre of tunnels experienced higher temperatures and reduced wind speed. In raspberry, fruit shape was improved with greater pollinator abundance and was susceptible to higher temperatures. In blueberry, per plant yield and mean berry weight were positively associated with pollinator abundance and were lower at the centre of tunnels than at the edge. Fruit quality (crumbliness) in raspberries was improved with a greater number of visits by sweat bees, who were not as susceptible to climatic conditions within tunnels. Understanding bee foraging behaviour and changes to yield under protected cropping conditions is critical to inform the appropriate design of polytunnels and aid pollinator management within them.

## 1. Introduction

With increasing worldwide demand for fresh fruit and vegetables, producers are increasingly turning to protected cropping in order to expand suitable growing regions, stabilize yields and improve product quality (Lamont, 2009). This involves housing the crop in a facility built with glass or semi-transparent plastic to protect it from external natural hazards, such as temperature extremes, heavy rain and wind damage (Gruda and Tanny, 2014; Lang, 2014). Such structures also allow for manipulation of the crop micro-environment to facilitate optimal plant growth, induce early flowering and extend fruit production duration, allowing growers to produce valuable out-of-season produce (Gruda and Tanny, 2014).

Plant physiological responses to protected cropping environments are comparatively well known (Gruda and Tanny, 2014), yet few studies have focused on the interactions between plant responses and pollinator performance within enclosures (Ariza et al., 2012; Dag and Eisikowitch, 1999). Pollinator-dependent crops grown in protected environments, such blueberry (*Vaccinium* spp.) and raspberry (*Rubus idaeus* L.), require additional management considerations given they rely on pollinator performance to optimize yield and quality (Andrikopoulos and Cane, 2018; Benjamin and Winfree, 2014; Klein et al., 2007). Yet, despite increasing adoption of protected cropping practices worldwide, little is known about the behavior, performance and response to climatic conditions of beneficial insect pollinators in these systems.

Unventilated protected cropping environments commonly have higher temperature and humidity and lower radiation levels than the external environment (Harmanto et al., 2006). Wind speed is reduced and plants are provided with consistent water, nutrients and in some systems, CO_2_ (Gruda and Tanny, 2014). The capacity to control these factors may act to enhance growing conditions for some crop species (Bakker, 1989; Mortensen, 1987; Slack and Hand, 1983; Wittwer and Robb, 1964). For instance, the environmental conditions under protected cropping conditions can alter flower abundance and longevity, nectar volume and concentration, and pollen dehiscence compared to plants grown in open environments (Corbet et al., 1979; Dag and Eisikowitch, 1999). Such changes to plant physiology likely influence bee foraging behaviour and longevity in protected cropping systems through altered resource availability (Di Pasquale et al., 2013; Vaudo et al., 2015). The environmental conditions themselves under covers may further limit pollinator health, activity and foraging efficiency through more direct means (Nielsen et al., 2017; Evans et al., 2019).

Most commercial raspberry and blueberry cultivars benefit from insect pollination to obtain higher quality fruit (Benjamin and Winfree, 2014; Ehlenfeldt, 2001; Ehlenfeldt and Martin, 2010; Zurawicz, 2016). Here, we used raspberry and blueberry as model crops to investigate bee activity on flowers and changes to fruit yield and quality along the length of polytunnels. This study had four main aims:

1. To determine how pollinator visitation rates vary with distance from the edge of commercial (∼100 m) polytunnels
2. To investigate whether microclimatic environmental conditions (e.g. temperature and relative humidity) vary along the length of polytunnels
3. To investigate the extent to which fruit yield and quality vary along polytunnels
4. To understand which explanatory variables best explain how fruit yield and/or quality vary along polytunnel lengths

## 2. Methods

### 2.1 Site selection and study design

Northern NSW and Far North Queensland are major regions for the protected cultivation of both crops, which are principally pollinated by both managed and wild bees, including a species of Australian native stingless bee *(Tetragonula carbonaria* Smith). Most farms stock European honeybee (*Apis mellifera* L.) colonies for pollination services, but wild stingless and solitary bees can provide a free crop pollination service and the commercial use of stingless bees is also growing in Australia (Halcroft et al., 2013; Heard and Dollin, 2000).

#### Raspberry

The raspberry study site was located in Corindi, New South Wales (29° 59’17”S, 153°08’27”E, Fig. 1a). The farm is a commercial enterprise of ∼ 325 ha managed by the Costa group, adopting conventional agricultural practices to the production of raspberry cultivars on approximately 95 ha and blueberry cultivars on 240 ha. Raspberry was grown in single variety blocks under ∼ 100 m long polythene-protected tunnels (polytunnels) at both east-west and north-south orientations (Fig. 1b). Each tunnel covered three rows of plants separated by 2.85 m. Domesticated European honeybees were stocked at a rate of 8 hives ha^-1^ across the farm, whilst native stingless and other bees were from wild populations nesting in surrounding forest and bare earth in and around the farm.

**Figure 1:**
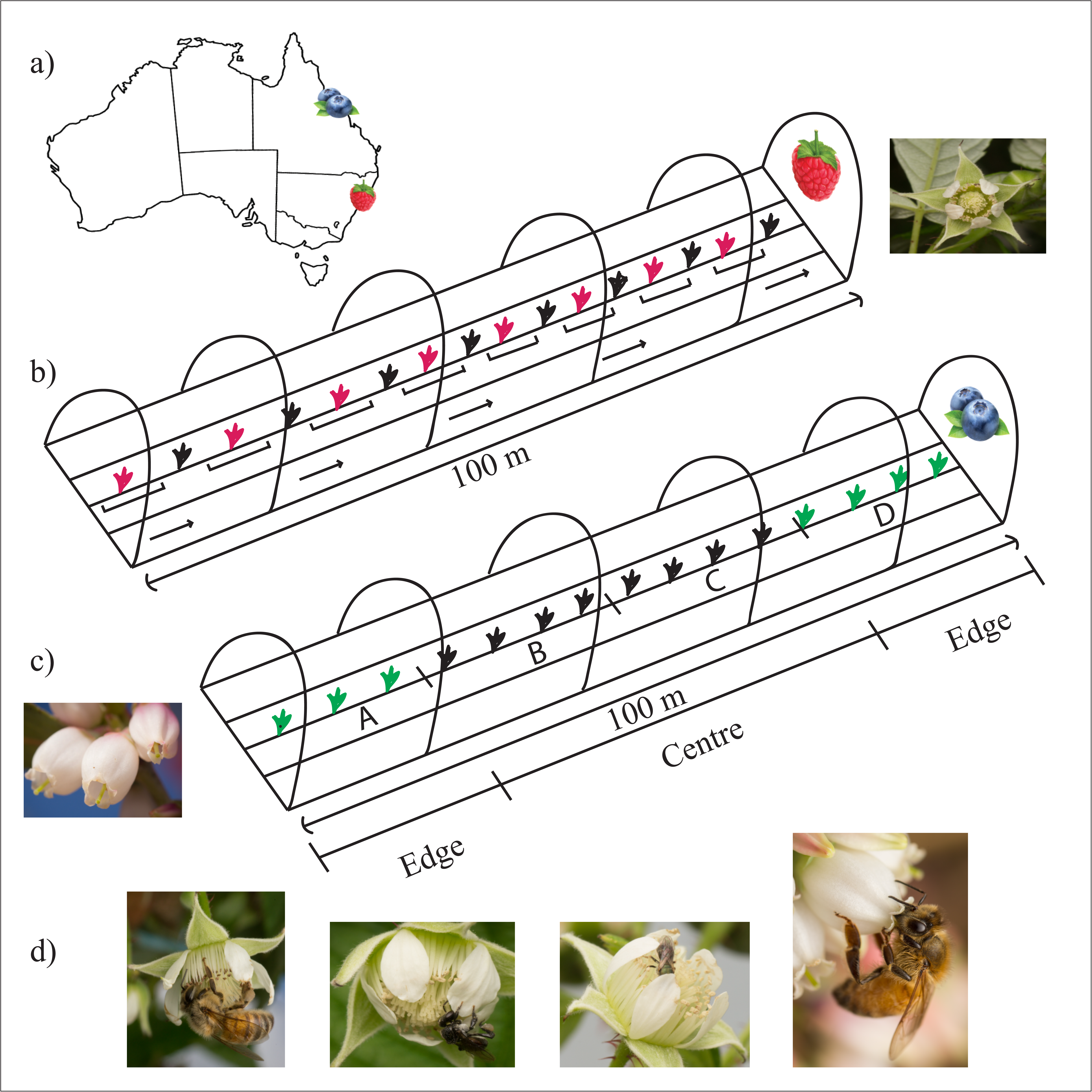
a) Location of raspberry farm in Corindi, NSW and blueberry farm in Walkamin, QLD, Australia; b) representation of raspberry polytunnels, where every fifth plant was surveyed along the row and fruit collected from these bushes to determine fruit quality.; c) representation of blueberry polytunnels and fruit harvesting segments. Also included are pictures of typical flowers and fruits. Centre rows of each surveyed tunnel were partitioned into four subsections A-D. Fruit collected from bushes located in subsections B and C were combined for total ‘centre’ yield and subsections A and D were combined for ‘edge’ yield; and d) Bee visitors to both raspberry and blueberry flowers, showing different foraging techniques used based on flower morphology.

#### Blueberry

The blueberry study site was located in Walkamin, Queensland (17° 06’49”S, 145°23’42”E, Fig. 1a). This site was a 15 ha commercial blueberry farm, also managed by the Costa group, utilising conventional agricultural methods. Evergreen highbush blueberry, *Vaccinium corymbosum* L. hybrids, were grown in single variety blocks under east-west orientated, high-set, 100 m long polytunnels (Fig. 1c). Each tunnel covered three rows of plants separated by 2.85 m. Blueberry plants were pruned to encourage flowering on primocane growth. Managed honeybees were stocked at a rate of 5 hives ha^-1^ across the farm. In addition to honey bees, managed hives of stingless bees were positioned adjacent to tunnels later used for pollinator surveys. These hives were positioned 3 m away from tunnel openings at the eastern side, one in front of every second tunnel, at a density of 11.7 hives ha^-1^. Wild colonies of *Tetragonula* sp. were also present around the farm.

Due to variation in the number of available polytunnels between the two study locations and differences in plant growth and flower morphology between the berry species, pollinator surveys and fruit collection were conducted in slightly different but complementary ways at the two sites.

### 2.2 Pollinator surveys

#### Raspberry

Pollinator surveys were conducted in the Austral late spring, summer and early autumn, between November 2017 and March 2018. Suitable survey tunnels were selected based on the flowering period of raspberry plants and the presence of stingless bees within tunnels. To reduce edge effects, the middle row (of three within each tunnel) was selected for surveys (n=23). Starting with the plant closest to the edge of the tunnel, surveys were conducted within a defined area – from the middle of the plant in the pot to half way between it and the pot to either side (Fig. 1b). Surveys were conducted for one minute, recording the identity of all individuals visiting the reproductive parts of flowers (e.g. honeybee, stingless bee). This was then repeated at every fifth plant along the row to the opposite end of the tunnel, giving a spacing of ∼ 4 m between survey points. In March 2018, surveys were conducted at a subset of tunnels (n=8, with two tunnels at a time surveyed consecutively over a three-day period) across three different time-periods (10:00-12:00, 12:00-14:00 and 14:00-16:00). This ensured that each tunnel was surveyed across each of the three time periods on different days. For instance, on day one a survey was conducted during 10:00-12:00 at one tunnel and during 12:00-14:00 in the other, then rotated through the other time periods on subsequent days. Remaining pairs of tunnels were then surveyed following the same method.

#### Blueberry

Transects were conducted down the entire 100 m length of the centre row of each of nine tunnels. Six transects were conducted per day in each tunnel at hourly intervals between 10:00 and 15:00, with each tunnel being visited for a total of three randomised days during fine weather. Each hourly transect was conducted on the opposite side of the row to the previous transect, for a total of three transects on the northern side of the row and three transects on the southern side of the row, per day. Transects were walked at a slow pace (12 m min^-1^) with observers scanning flowers for insect visitors on the closest half of the plant from the centre of the row. Transects were walked into the direction of the Sun to avoid shadows disturbing foraging insects. Tallies of insect flower-visitors were recoded for each 4 m length of row, encompassing five blueberry plants (Fig. 1c). The site was visited during the peak blooming period, between 31 April – 17 May 2018.

### 2.3 Microclimate data

#### Raspberry

Microclimatic conditions within tunnels were recorded during the period of consecutive three-day surveys in March 2018. Twelve Kestrel DROP 2 temperature and relative humidity (RH) loggers (© Nielsen-Kellerman) were placed in each of the two survey tunnels, spaced evenly along the length of the tunnel. These remained in place for three days to record environmental conditions whilst repeat insect surveys were performed, then rotated to other tunnels to repeat the process—for a total of eight tunnels over a two-week period. Loggers were hung above the raspberry plants and attached with cable ties (Fig. 1d). The average wind speed at each survey point along the tunnel was also recorded during insect surveys using a Kestrel 2000 wind meter (© Nielsen-Kellerman). This was held so it faced along the tunnel, recording the average wind speed after ten seconds. Light intensity, both UV and white light, were recorded at each survey point, using a LumaCheck Dual UV and White Light Meter (© Baugh and Weedan Ltd.).

#### Blueberry

Microclimatic conditions within tunnels were recorded daily during the period of transect walks, conducted between March and April 2018. For each of the two tunnels surveyed per day, a group of three to five Kestrel DROP 2 temperature and relative humidity loggers were positioned equally along the length of the survey tunnels. Loggers remained in place for a twenty-four-hour period to record environmental conditions whilst repeat insect surveys were performed. Loggers were positioned along the centre row of plants at a height of 1.2 metres above ground and were protected from solar radiation by two layers of inverted white polypropylene plastic bowls, suspended 10 cm above the temperature sensor (Fig. 1d). At the conclusion of each day, the two sets of loggers were positioned into the following day’s survey tunnels, in a randomised rotating system.

### 2.4 Fruit yield and quality

#### Raspberry

In March 2018, following completion of the consecutive tunnel visits, one stem (containing at least five flowers/early fruits) on each of the survey plants (every fifth plant) for the first half of the transect, were bagged using white mesh organza bags. When berries ripened, bags were removed, fruits picked and measurements taken—height, width and fruit weight, the number of druplets and fruit crumbliness (an industry measure of quality, (Daubeny et al., 1967; Jennings, 1967).

#### Blueberry

The surveyed centre-row of each tunnel was divided into four sections along the entire length of the tunnel. These subsections were demarcated with flagging tape and paint.

The number of plants in each subsection was recorded. Berries were harvested from every surveyed row by hand at approximately ten-day intervals over the course of the entire harvest period which occurs 2-3 months after flowering, between May and August 2018. For each harvesting event, the total berry yield and the combined weight of twenty-five randomly sampled berries for each subsection was recorded. Data from these four sections was pooled into two groups representing edge and centre sections (Fig. 1c).

### 2.5 Statistical analyses

All statistical analyses were conducted in R (v.3.5.1, R Core Team, 2018). We conducted five separate analyses to address our study aims. First, to test if each of the microclimatic factors recorded (raspberry: average temperature, relative humidity, average wind speed, UV light and white light; blueberry: average temperature, relative humidity) varied with increasing distance into polytunnels, we used univariate generalised linear models using the *lme4* package (Bates et al., 2014), assuming a Poisson distribution. Where overdispersion was detected (raspberry: relative humidity, wind speed, UV and white light models; blueberry: relative humidity model) a negative binomial distribution was set (Zuur et al., 2009). Second, to test the influence of distance from edge and microclimatic conditions on bee foraging within polytunnels, we used generalised linear mixed models using the *lme4* package (Bates et al., 2014), with a negative binomial distribution and logit-link function. Separate models were run for each of the three bee species most commonly recorded: honeybees, stingless bees and a species of sweat bee (*Homalictus urbanus* Smith). For blueberry, only honeybees were recorded in sufficient number to be included in the analysis. A nested random factor of *block and row* by *survey date* was included in models to account for resampling within tunnels. Two variables (relative humidity and white light) were highly correlated (>0.7) with other variables and were removed from models prior to analysis. As values differed by orders of magnitude, all predictor variables were rescaled prior to analyses, using the *scale* function. Third, we tested whether fruit yield and quality differed between the edge and centre of tunnels using Welch’s t-tests. For raspberry, we tested whether characteristics of fruit quality (weight, height, width, number of druplets and crumbliness) varied with position in polytunnels (edge versus centre). Fruits from all surveyed plants within each section were grouped. For blueberry, average berry weight and total yield per plant were used as response variables. Fourth, to assess if these fruit quality metrics correlated with the abundance of honeybees, stingless bees and total bee abundance in raspberry, and honeybee abundance in blueberry, we performed separate Spearman rank correlations for each yield and quality metric. Finally, to test which of our explanatory variables (visitation rates or microclimate) best explained differences in fruit yield and/or quality, we used generalised linear mixed models using the *glmmadmb* package (Skaug et al., 2011), with a negative binomial distribution and logit-link function. Again, separate models were run for each of the three bee species most commonly recorded: honeybees, stingless bees and sweat bee, as well as for total visitation of all species. For blueberry, only honeybees were recorded in sufficient number to be included in the analysis. A nested random factor of *block and row* was included in models to account for resampling within tunnels.

## 3. Results

### 3.1 Flower visitation and bee abundance along polytunnel lengths

We recorded a total of 5698 honeybee (*A. mellifera*), 2034 stingless bee (*T. carbonaria*) and 250 sweat bee (*H. urbanus*) visits to raspberry flowers (n=1054 survey points across 23 polytunnels) and 1762 honeybee visits to blueberry flowers (n=4411 survey points across nine polytunnels). Other insects recorded in very low numbers foraging in blueberry tunnels included hover flies (Syrphidae, n=54), house flies (Muscidae) and butterflies (Lepidoptera) (combined n=25). Only 15 stingless bees and no sweat bees were recorded foraging on blueberries across the study, and thus were not included in analyses for this crop.

#### Raspberry

We found a greater number of raspberry flower visits by honeybees (Est=-0.17, SE=0.06, P<0.01) and stingless bees (Est=-1.29, SE=0.09, P<0.01) per minute at the ends of tunnels, and fewer visits to flowers toward the middle of tunnels (Fig. 2). This led to a reduction in visits per minute, from ∼7 visits per minute down to ∼5.5 visits for honeybees and from ∼4.5 visits per minute to ∼0.5 visits for stingless bees (Fig. 2a). The wild native bee species *H. urbanus* (Halictidae) provided lower but consistent visitation throughout the length of tunnels (Est=-0.12, SE=0.19, P=0.53), relative to honeybees and stingless bees (Fig. 2a).

**Figure 2:**
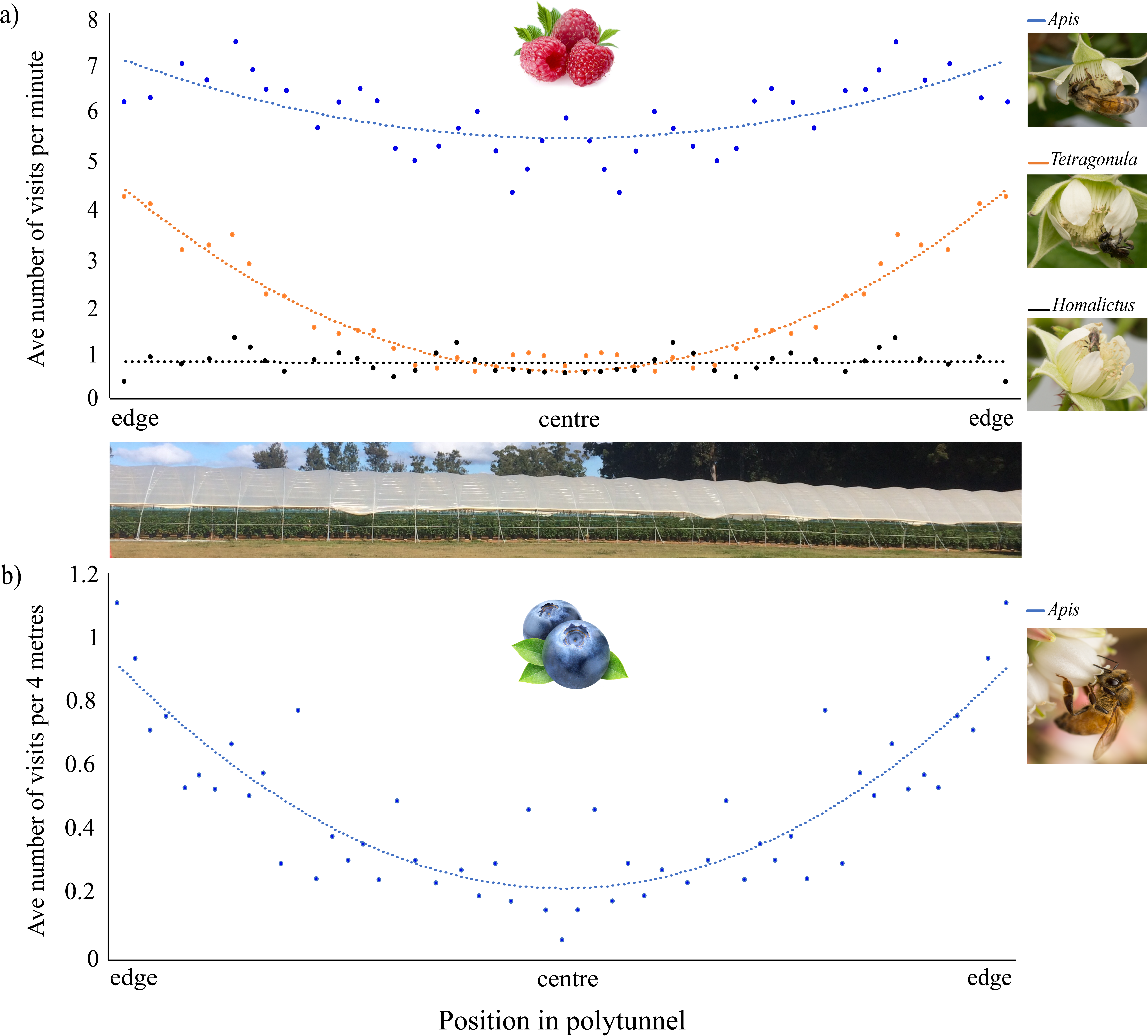
a) The average number of visits per minute to raspberries and b) the number of visits per 4 m section of tunnel (at a walk rate of ∼ 14 m per min) to blueberries by honeybees *Apis mellifera*, stingless bees *Tetragonula carbonaria* and sweat bees *Homalictus urbanus* along polytunnels used for commercial crop production (pictured).

#### Blueberry

As with raspberry, we found a greater number of visitors to blueberry flowers by honeybees (Est=-1.03, SE=0.08, P<0.01) per area search at the ends of tunnels, and fewer visitors to flowers toward the middle of tunnels. This led to a reduction in visitor abundance, from an average of ∼0.9 visitors per five blueberry plants at the ends of tunnels to ∼0.2 visitors per five blueberry plants in the centre of tunnels (Fig. 2b).

### 3.2 Microclimatic conditions along polytunnel lengths

Microclimatic conditions changed along the length of polytunnels (Fig. S1). Average temperature increased with increasing distance into tunnels from the edge (raspberry: Est=0.001, SE=0.0005, P<0.01, Fig. S1a; blueberry: Est=0.0004, SE=0.0002, P=0.03, Fig. S1d), whilst relative humidity decreased for raspberry (Est=-0.002, SE=0.0008, P<0.01, Fig. S1b), but remained fairly constant for blueberry (Est=-0.0004, SE=0.0002, P=0.08). Average wind speeds also decreased further into polytunnels containing raspberry (Est=-0.03, SE=0.006, P<0.01, Fig. S1c), whilst both UV and white light remained unchanged throughout tunnels (UV: Est=0.002, SE=0.003, P=0.38; white light: Est=0.006, SE=0.004, P=0.16).

#### 3.2.1 Microclimate variables in relation to pollinator visitation and abundance

##### Raspberry

Visitation by honeybees, stingless bees and sweat bees increased with increasing average temperature (honeybee: Est=0.39, SE=0.09, P<0.01, Fig. 3a; stingless bee: Est=0.97, SE=0.16, P<0.01, Fig. 3b; sweat bee: Est=0.92, SE=0.39, P=0.02, Fig. 3c). Both honeybee and stingless bee visits also increased with increasing wind speed inside the polytunnel (honeybee: Est=0.16, SE=0.06, P=0.01, Fig. 3d; stingless bee: Est=0.21, SE=0.09, P=0.02, Fig. 3e). UV light had no significant effect on visitation.

**Figure 3:**
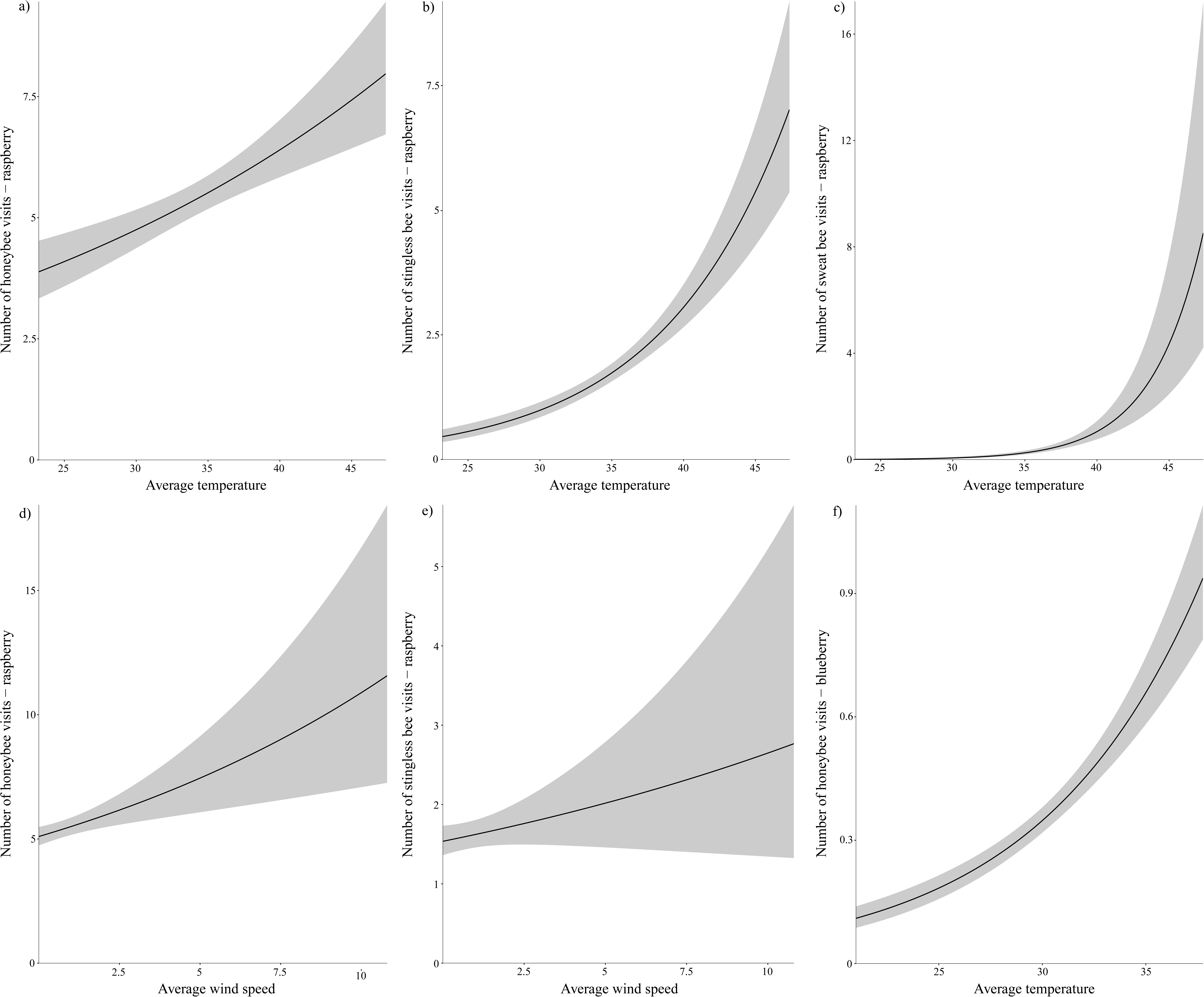
The response of bee species to microclimatic conditions within polytunnels: the effects of increasing temperature on the number of visits to raspberries by a) honeybees, b) stingless bees and c) sweat bees; the effects of increasing wind speed on the number of visits to raspberries by d) honeybees and e) stingless bees; the effects of increasing temperature on the number of visits to blueberries by f) honeybees.

##### Blueberry

Honeybee abundance increased with increasing temperature inside the polytunnel (Est=0.62, SE=0.11, P<0.01, Fig. 3f).

### 3.3 Yield and fruit quality along polytunnel lengths

#### Raspberry

Of all five fruit quality measures, none showed any significant difference between the edge and centre of tunnels (P > 0.05). However, there was a significant weak positive relationship between stingless bee abundance and both fruit height (ρ = 0.214) and number of raspberry druplets (ρ = 0.260) and between all bee abundance and the number of druplets (ρ = 0.206) across all polytunnels (Spearman’s correlation test: stingless bee abundance ∼ average fruit height, S = 165, p-value = 0.026; stingless bee abundance ∼ average number of druplets, S = 155, p-value = 0.007; all bee abundance ∼ average number of druplets, S = 167, p-value = 0.033, Figs. 5a-5c).

**Figure 4.**
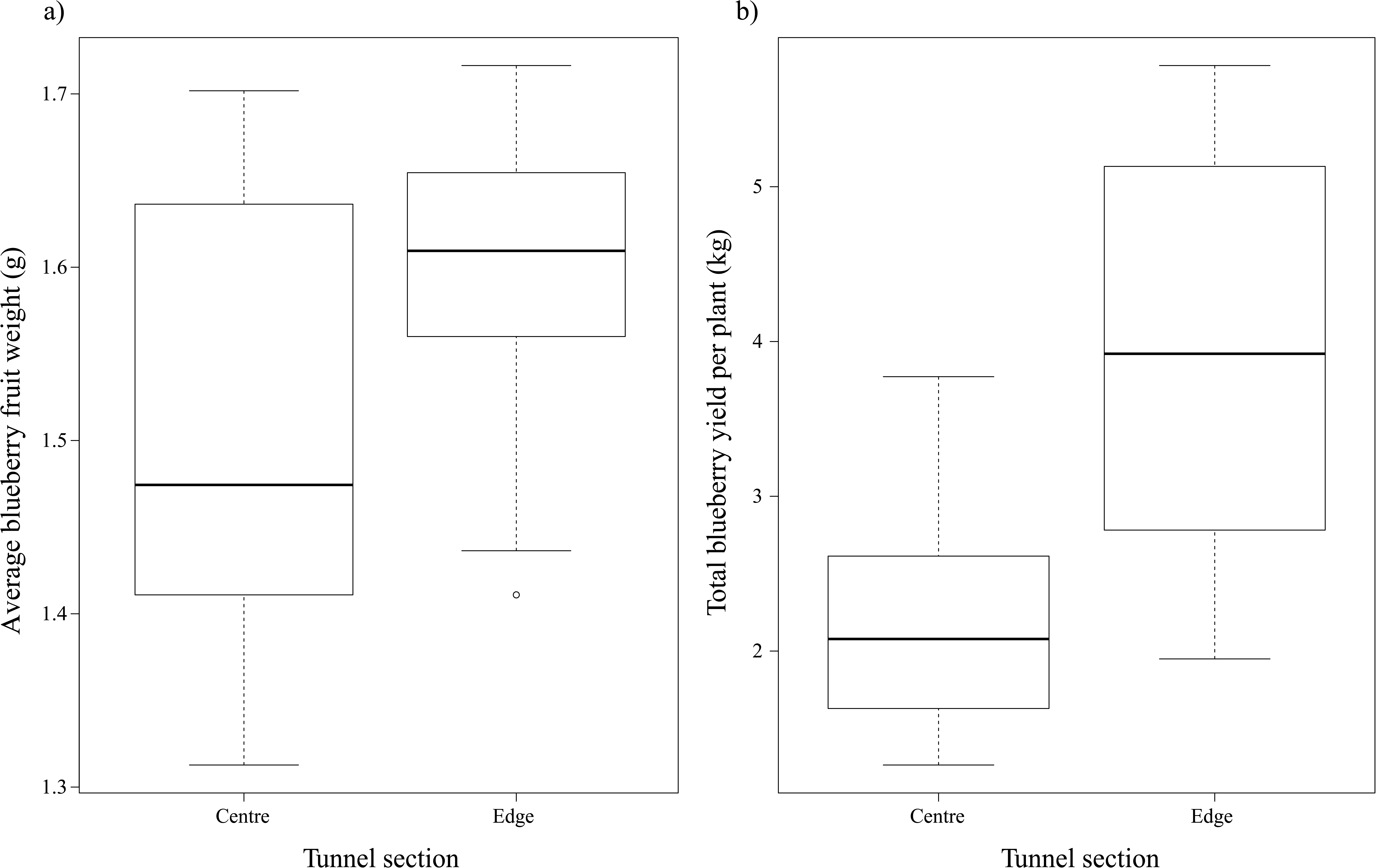
a) Average blueberry fruit weight (grams) and b) total blueberry yields per plant (kg) at the centre and edges of polytunnels.

**Figure 5.**
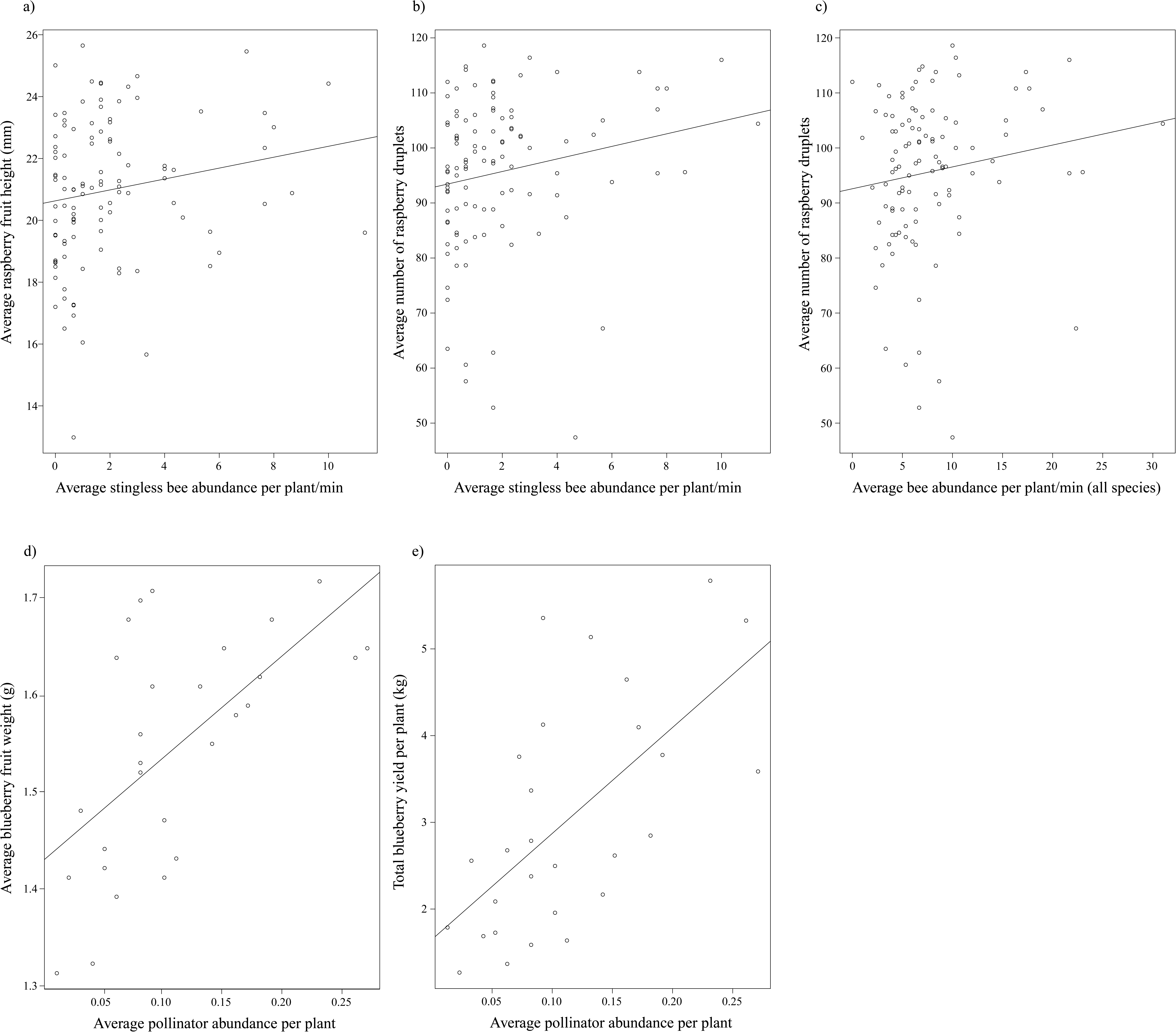
a) Relationship between average abundance of stingless bees per plant/min and average raspberry height (mm), b) average abundance of stingless bees per plant/min and average number of raspberry druplets, c), average abundance of all bees per plant/min and average number of raspberry druplets, d) average abundance of bees per plant and associated average blueberry weight (g), and e) average abundance of bees per plant and associated total blueberry yield.

#### Blueberry

Average berry weight and average yield per plant was significantly higher for plants at the edges of tunnels compared to the centre (Welch’s t-test: average berry weight, t = −2.43, df = 23.2, P < 0.05; total yield per plant, t = −4.01, df = 20.8, P < 0.001, Figs. 4a, 4b). There were also significant positive relationships between pollinator abundance and average blueberry weight (ρ = 0.605) and yield per plant (ρ = 0.633) across the different sections of the polytunnels (Spearman’s correlation test: berry weight, S = 1443.4, n = 28, P < 0.001; yield per plant, S = 1342, n = 28, P < 0.001, Figs. 5d, 5e).

### 3.4 Which variables best explain variation in fruit yield and quality?

#### Raspberry

Fruit shape (height and width) and number of druplets were most reduced by increased temperature conditions (Table 1). Fruit quality (crumbliness) was improved with greater visitation by sweat bees. Fruit crumbliness was also negatively affected by the interaction between this bee species and average temperatures along the length of polytunnels (Table 1).

**Table 1:** Estimated contribution of bee visitation and average temperature to each of the fruit yield and quality measures in raspberry and blueberry. Significant responses in bold.

| Model |  | Visitation |  |  | Ave. Temp |  |  | Interaction |  |  |
| --- | --- | --- | --- | --- | --- | --- | --- | --- | --- | --- |
|  |  | Est | SE | Pr(>\|z\|) | Est | SE | Pr(>\|z\|) | Est | SE | Pr(>\|z\|) |
| *Raspberry* |  |  |  |  |  |  |  |  |  |  |
| All visitors | Fruitshape ~ All * Ave Temp | -0.02 | 0.04 | 0.54 | **-0.02** | **0.01** | **0.04** | 0.00 | 0.00 | 0.48 |
|  | Crumbliness ~ All * Ave Temp | -0.01 | 0.03 | 0.71 | -0.01 | 0.01 | 0.36 | 0.00 | 0.00 | 0.71 |
|  | Druplets ~ All * Ave Temp | 0.02 | 0.05 | 0.70 | -0.01 | 0.01 | 0.23 | 0.00 | 0.00 | 0.69 |
|  | Weight ~ All * Ave Temp | 0.00 | 0.05 | 0.95 | 0.00 | 0.01 | 0.86 | 0.00 | 0.00 | 0.95 |
| Honeybee | Fruitshape ~ HB * Ave Temp | -0.02 | 0.05 | 0.68 | -0.02 | 0.01 | 0.09 | 0.00 | 0.00 | 0.70 |
|  | Crumbliness ~ HB * Ave Temp | -0.01 | 0.04 | 0.73 | -0.01 | 0.01 | 0.39 | 0.00 | 0.00 | 0.76 |
|  | Druplets ~ HB * Ave Temp | 0.04 | 0.07 | 0.60 | -0.01 | 0.01 | 0.23 | -0.00 | 0.00 | 0.67 |
|  | Weight ~ HB * Ave Temp | 0.01 | 0.07 | 0.88 | 0.00 | 0.01 | 0.73 | 0.00 | 0.00 | 0.86 |
| Stingless bee | Fruitshape ~ SB * Ave Temp | -0.07 | 0.09 | 0.48 | **-0.02** | **0.01** | **0.03** | 0.00 | 0.00 | 0.35 |
|  | Crumbliness ~ SB * Ave Temp | -0.05 | 0.07 | 0.49 | -0.01 | 0.01 | 0.25 | 0.00 | 0.00 | 0.44 |
|  | Druplets ~ SB * Ave Temp | -0.07 | 0.12 | 0.56 | **-0.02** | **0.01** | **0.03** | 0.00 | 0.00 | 0.62 |
|  | Weight ~ SB * Ave Temp | -0.08 | 0.12 | 0.52 | -0.00 | 0.01 | 0.90 | 0.00 | 0.00 | 0.48 |
| Sweat bee | Fruitshape ~ HU * Ave Temp | 0.63 | 0.46 | 0.17 | -0.01 | 0.01 | 0.15 | -0.02 | 0.01 | 0.16 |
|  | Crumbliness ~ HU * Ave Temp | **0.82** | **0.41** | **0.04** | 0.00 | 0.01 | 0.76 | **-0.02** | **0.01** | **0.03** |
|  | Druplets ~ HU * Ave Temp | -0.31 | 0.62 | 0.61 | -0.02 | 0.01 | 0.05 | 0.01 | 0.02 | 0.69 |
|  | Weight ~ HU * Ave Temp | 0.83 | 0.58 | 0.15 | 0.01 | 0.01 | 0.46 | -0.02 | 0.02 | 0.14 |
| *Blueberry* |  |  |  |  |  |  |  |  |  |  |
| Honeybee | Weight ~ HB * Ave Temp | **0.08** | **0.02** | **<0.01** | -0.02 | 0.02 | 0.24 | 0.00 | 0.020 | 0.96 |
|  | Total yield ~ HB * Ave Temp | **1.08** | **0.18** | **<0.01** | **-0.70** | **0.19** | **<0.01** | -0.45 | 0.24 | 0.06 |

#### Blueberry

Average berry weight and total berry yield per plant both increased with greater visitation by honeybees (Table 1). Total yield per plant was also reduced with increased temperature conditions (Table 1).

## 4. Discussion

The results of this study demonstrate that polytunnels vary in conditions for fruit production along their length. First, we found that forager activity was greatly reduced in the centre of tunnels for both crops. This result supports similar reduced honeybee visitation to raspberry grown under protected cropping in Denmark (Nielsen et al., 2017). The close proximity of tunnels to both managed hives and native vegetation that supports wild bees is such that we would not expect bee foraging to differ across the length of tunnels under optimal conditions. Plant age and flowering density were consistent throughout tunnels, and both honeybees and stingless bees are known to travel several hundred metres to several kilometres to locate food sources (Beekman and Ratnieks, 2000; Smith et al., 2017). Thus, a distance of ∼100 m would be easily navigable by foragers unless other factors prevented them from effectively reaching the centre of tunnels.

Second, temperature and humidity were higher in the centre of tunnels, relative to the edges, yet wind speed was lower. While increasing temperatures are generally associated with the onset of bee foraging activity (Corbet et al., 1993; Heard and Hendrikz, 1993; Tan et al., 2012), the drop in activity in the centre of tunnels was likely due to a range of interacting factors. Several managed bee species are susceptible to thermal extremes (heat or cold) which impact their effectiveness in enclosed environments (da Silva et al., 2017; Kovac et al., 2014). Conditions inside protected cropping environments are frequently warmer and prolonged exposure to high temperatures may have lethal or sub-lethal effects on some bee species (da Silva et al., 2017). For instance, foraging European honeybees exposed to prolonged temperatures above 40°C have reduced lifespans, whilst elevated hive temperatures during pupal development can impact wing formation, adult learning and memory (Abou-Shaara et al., 2017, 2012; Tautz et al., 2003). Highly humid conditions further diminish the ability of bees to thermo-regulate, resulting in reduced tolerance to high temperatures which may occur in tunnel centres (Free and Spencer-Booth, 1962).

Other microclimatic conditions associated with protected cropping environments may also directly impact bee foraging behaviour, colony dynamics and health (Lang, 2014). For instance, as bees are highly sensitive to ultraviolet (UV) light (Peitsch et al., 1992), polythene coverings used in commercial polytunnels alter light intensity and impact bee foraging (Morandin et al., 2001). If conditions are not ideal for bee foraging, colonies may suffer physical stress, disease or poor nutrition, or may simply avoid foraging in these environments (Dag, 2008; Pinzauti, 1994; Whittington and Winston, 2003). Whilst UV light did not change across the length of tunnels here, the effect of altered light may be intensified as bees forage further into tunnels, particularly when they are already under heat stress, limiting their ability to successfully access flowers toward the centre.

Third, we found that average berry weight and overall fruit yield in blueberries and quality in raspberries, were reduced in the centre of polytunnels. Changes in yield down the length of tunnels was negatively correlated with bee abundance, indicating pollinator foraging activity is likely related to high-quality fruit, as in other studies (Garratt et al., 2014; Klatt et al., 2014; Malagodi-Braga and Kleinert, 2004; Slaa et al., 2006). Fruit quality is an important commercial consideration in raspberry, as under-sized or misshapen fruits affect their marketability (Ariza et al., 2012; Wietzke et al., 2018). In this study, altered fruit shape and number of druplets was recorded in raspberries along the length of tunnels. Fruit quality (crumbliness) however was improved by greater visits by the wild native sweat bee, similar to findings in strawberry crops (MacInnis and Forrest, 2019). Visits by this species may have offset some of the loss of quality incurred by reduced visits by both honeybees and stingless bees at the centre of tunnels. Thus, the presence of sweat bees in some tunnels likely made fruits more marketable, ultimately improving profit to farmers.

Finally, we found that microclimatic conditions in tunnels had a more direct impact on fruit quality than bee visitation in raspberry, while in blueberries, increased honeybee visitation improved both yield and average weight of berries, but higher temperatures reduced total blueberry yield. This indicates that increased temperature at the centre of tunnels is likely driving the responses here, both to the number of visits to flowers by pollinators, and the resulting fruit set in berry crops. In greenhouse-grown tomato, high temperatures are known to negatively affect fruit-set, pollen viability and pollen release, which may be improved by lowering temperature and increasing humidity (Sato et al., 2006; Harel et al., 2014). Within cropping enclosures, temperature and relative humidity interact to alter vapour pressure deficit (VPD), which in turn affects plant respiration and disease susceptibility (Gruda & Tanney, 2014). How animal-pollinated crops respond to heat stress and altered VPD in protected cropping conditions still remains largely unexplored. If pollinator abundance and berry yields are to be improved, optimal conditions need to be maintained across the length of tunnels, probably through physical alteration to current structures to reduce heat stress and improve airflow.

Understanding the environmental and management conditions that impact pollinator dependent crop yields are imperative in order to optimize yields in protected cropping systems. Studies to date have identified a number of pollinator species that forage effectively under enclosed conditions, including honeybees, stingless bees, hoverflies (Diptera: Syrphidae) and bumblebees (*Bombus* spp.) (Malagodi-Braga and Kleinert, 2004; Slaa et al., 2006; Trillo et al., 2018). Given the abundance of stingless bees and the biosecurity and biodiversity risks concerns of introducing bumblebees to mainland Australia (Dafni et al., 2010; Kingston and McQuillan, 1998), stingless bees provide the best safeguard against potential future pollination deficiencies (Allsopp, 2004; De la Rúa et al., 2009; Seitz et al., 2015). However, to effectively use stingless bees for crop pollination, optimal conditions must be maintained across the length of polytunnels. In addition to *T. carbonaria* and *A. mellifera*, this study also identified a sweat bee, *H. urbanus* foraging in some of the tunnels. However, little is known of the capacity of this species to pollinate other crops or their activity patterns in protected cropping conditions. *Homalictus* species nest in bare earth, often in aggregations — that is, many nests all clustered together, or many individuals using the same nest entrance (Michener, 1960). Given this nesting behaviour, there is potential for simple management strategies to attract this free wild pollinator to nest at the base of crops and provide a valuable contribution to pollination. Research and development on these and other wild and managed taxa that are already providing such pollination services is urgently needed to maximise our understanding of their biological needs (Cunningham et al., 2002; Jaffé et al., 2015). Such understanding will aid selection of the most appropriate polytunnel design to improve foraging conditions and maximize this vital pollination service to crop production.

## Acknowledgements

This work was funded by an Australian Research Council Discovery Early Career Researcher Award DE170101349 to Romina Rader and conducted in conjunction with Costa Group. Thanks to Raymond Dempsey for assistance in blueberry crops.

