## Supplementary materials for "Bee visitation and fruit quality in berries under protected cropping vary along the length of polytunnels"

Supplementary material: **Microclimatic conditions impact bee visitation and fruit quality in berries under protected cropping**

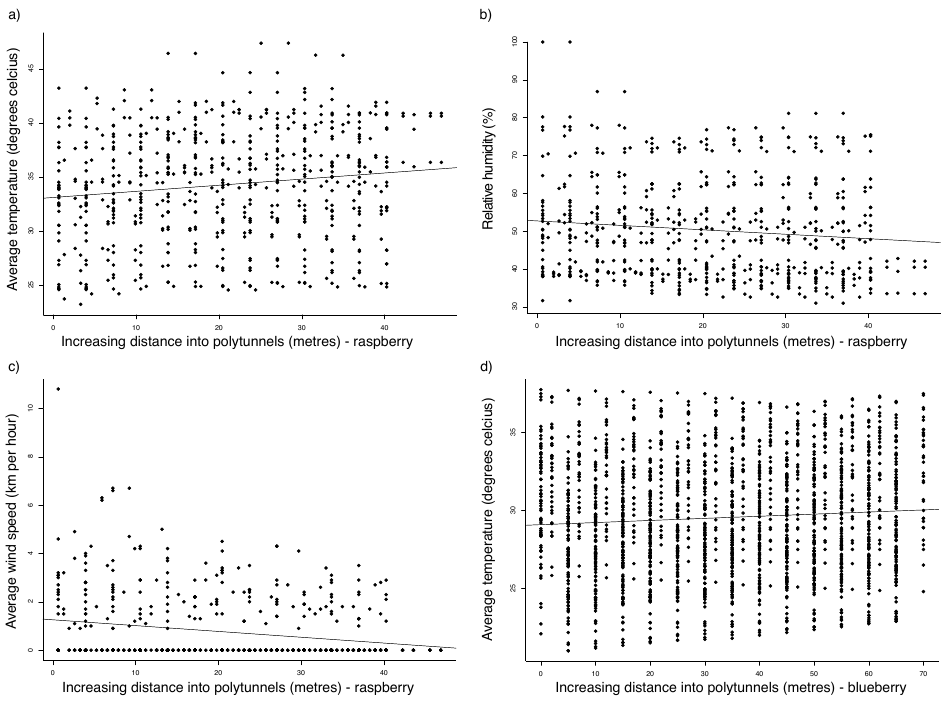

Figure S1: Changes in climatic condition with increasing distance into polytunnels used for raspberry and blueberry production (a) average temperature in raspberry polytunnels (b) relative humidity in raspberry polytunnels (c) average wind speed in raspberry polytunnels (d) average temperature in blueberry polytunnels.

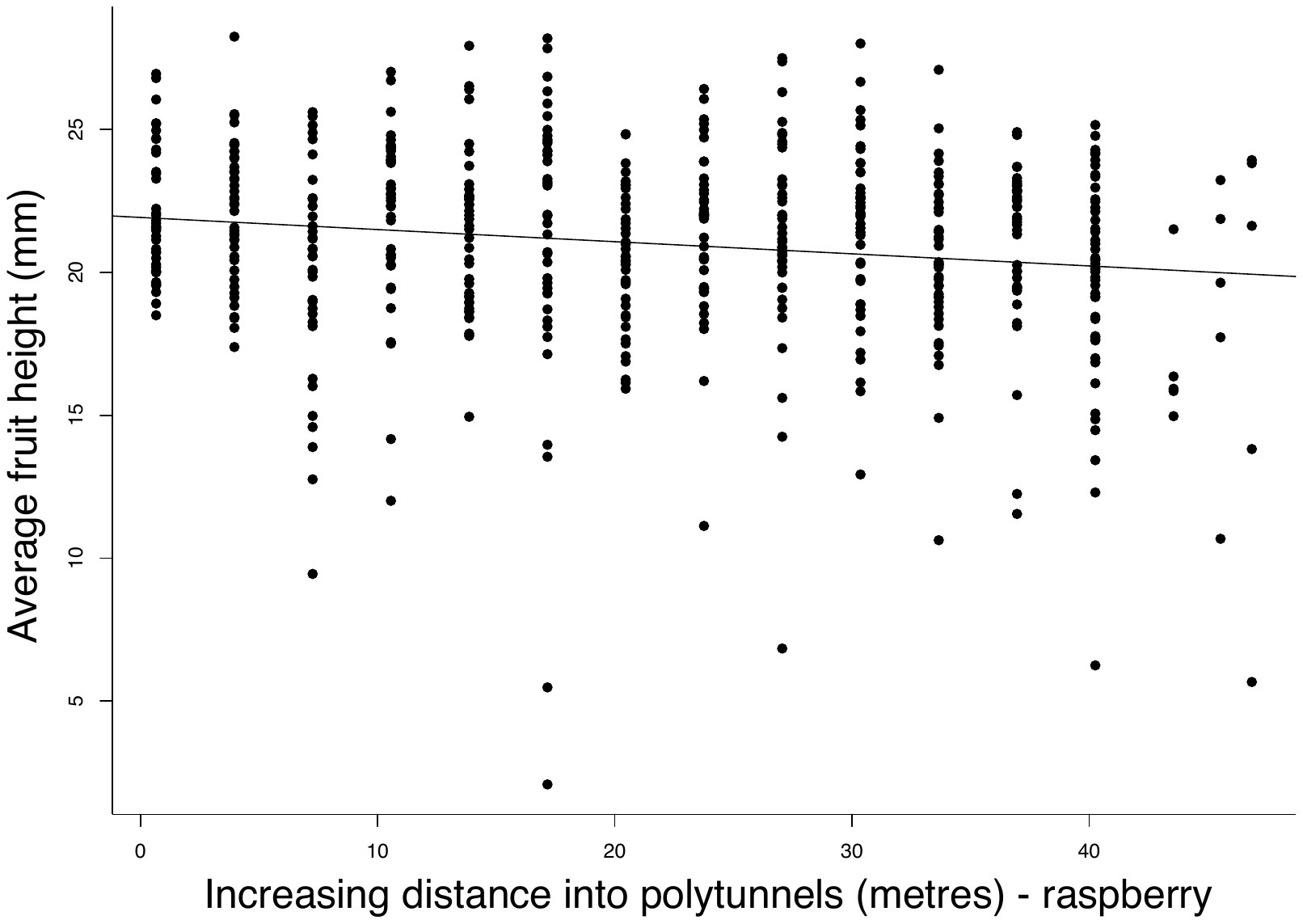

Figure S2: Variation in raspberry fruit height with increasing distance into polytunnels
